# Selective detection of m^6^A derived from mRNA using the Phospho-tag m^6^A assay

**DOI:** 10.1101/2022.05.23.493172

**Authors:** Aashiq H. Mirza, Nabeel Attarwala, Steven S. Gross, Qiuying Chen, Samie R. Jaffrey

## Abstract

N6-methyladenosine (m^6^A) is a modified nucleotide found in mRNA, ribosome RNA (rRNA) and small nuclear RNA (snRNA). m^6^A in mRNA has important roles in regulating mRNA stability, splicing, and other processes. Numerous studies have described m^6^A as a dynamic modification using mass spectrometry-based quantification of m^6^A in mRNA samples prepared from different cellular conditions. However, these results have been questioned based on the finding that the mRNA purification protocols often result in varying levels of rRNA contamination. Additionally, mRNA purification protocols disproportionately enrich for the 3’ ends of mRNA, a region that is enriched in m^6^A. To address these problems, we developed the Phospho-tag m6A assay, a highly efficient method for quantifying m6A specifically from mRNA. In this assay, a series of selective RNase digestion steps is performed, which results in m^6^A from rRNA and snRNA being liberated as m^6^A monophosphate, while m^6^A from mRNA is mostly liberated as m6A nucleoside. m^6^A levels are normalized to transcript levels, using m^7^G monophosphate liberated by yDcpS decapping enzyme as a surrogate for mRNA levels. Notably, this approach uses total cellular RNA, rather than purified mRNA, which simplifies the steps for m^6^A detection and overcomes the 3’-end biases associated with mRNA purification. Overall, the Phospho-tag m^6^A provides a simple and efficient method for quantification of mRNA-derived m^6^A from total RNA samples.

## Introduction

A major concept is that *N*^6^-methyladenosine (m^6^A), the most abundant internal nucleotide modification in mRNA, is “dynamic,” i.e., its levels can change in mRNA in different cell types or conditions. For example, changes in the levels of METTL3/METTL14, the biosynthetic heterodimeric methyltransferase that synthesizes m^6^A in mRNA^1,2^, are thought to lead to alterations in m^6^A levels in mRNA in specific disease states or cellular contexts^3^. These changes in m^6^A levels may affect the stability or processing of specific m^6^A-modified transcripts, thus affecting diverse cellular pathways and processes^4^.

The concept that m^6^A is dynamic comes from various types of m^6^A quantification assays, including dot blots and mass spectrometry, the latter of which is considered more quantitative. However, questions have been raised about the accuracy of these methods^5,6,7^. The major problem is that these methods require pure mRNA^8^. mRNA is typically <2%of total cellular RNA, while other m^6^A-containing RNAs such as snRNA, and especially rRNA, are ∼70% of total cellular RNA^9^. As a result, even highly efficient mRNA enrichment methods result in mRNA preparations that contain variable amounts of residual rRNA or snRNA^10^. For example, several studies demonstrated large amounts of residual rRNA levels even after two rounds of oligo-dT-based purification of mRNA^6^. rRNA-removal kits similarly leave considerable amounts of rRNA in samples^11^. Overall, these studies demonstrate that ostensibly pure mRNA fractions usually contain variable amounts of rRNA. Thus, putative differences in m^6^A in different samples could simply reflect differences in rRNA contamination.

A second major problem with using purified mRNA for m^6^A quantification is that mRNA is rarely purified as a full-length transcript. This is because RNA is highly susceptible to degradation by nucleases during purification. As a result, when mRNA is purified using oligo-dT resin, the resulting RNA fragments are biased for the 3’ end of the mRNA that contains the poly(A) tail^12^. This 3’ bias has been widely documented and seen to varying degrees in RNA-Seq studies^13^. Since m^6^A is enriched in mRNA 3’UTRs^14,15^, sample-to-sample variability in 3’ bias can result in different levels of m^6^A even if there are no underlying differences in methylation between samples. Overall, these highly prevalent artifacts of mRNA purification raise questions about the accuracy of previous m^6^A measurements which have lead to the conclusion that m^6^A is dynamic in the transcriptome.

To develop a method to reliably quantify m^6^A without the biases caused by mRNA purification, we developed the “phospho-tag” m^6^A assay. The phospho-tag m^6^A assay uses a simple and highly efficient sequential digestion protocol and relies on total cellular RNA rather than purified mRNA. The m^6^A that is liberated from the cellular RNA contains either a 5’-hydroxyl or a 5’-phosphate. m^6^A with a 5’-hydroxyl exclusively derives from mRNA, while m^6^A with a 5’-phosphate (i.e., m^6^AMP) derives from the alternative cellular m^6^A sources, i.e., rRNA and snRNA. Because these two forms of m^6^A have different masses, they can be readily resolved and quantified by mass spectrometry. In order to normalize mRNA-derived m^6^A levels, we developed a second approach. This method quantifies the total number of mRNA transcripts in a cell using a similar a phosphate-tagging method: m^7^G derived from mRNA caps contains a 5’-phosphate (i.e., m^7^GMP), while m^7^G derived from tRNA/rRNA contains a 5’ hydroxyl. The phospho-tag m^6^A assay is a “one-pot” reaction that simultaneously generates both m^6^A and m^7^GMP for quantification of m^6^A normalized to total mRNA levels. Together, the phospho-tag m^6^A assay provides a simple and rapid method for highly accurate quantification of mRNA-derived m^6^A from essentially any biological sample.

## RESULTS

### rRNA is highly prevalent in MS samples despite mRNA enrichment protocols

A potential problem with mass spectrometry-based m^6^A measurements is the possibility that rRNA is present as a contaminant in the poly(A) RNA fraction. mRNA purification using either oligo dT or rRNA depletion oligonucleotides has been shown to produce mRNA that can possibly have considerable levels of rRNA contamination^10,11^. If there is rRNA in these mRNA samples, this would be problematic since rRNA is known to contain m^6^A^16,17^ and could therefore account for the m^6^A detected by mass spectrometry. Notably rRNA contamination would lower the calculated m^6^A prevalence in a sample since m^6^A levels in rRNA is lower than m^6^A levels in mRNA. The m^6^A prevalence is 0.23% of adenosines in 18S rRNA and 0.23% in 28S rRNA (1 m^6^A in the 1869 nt 18S rRNA and 1 m^6^A in the 5070 nt 28S rRNA). In contrast, m^6^A prevalence in mRNA is approximately twice as high—the level of m^6^A in mRNA is reported to be 0.4% of adenosines^14,15^. Thus, variation in the degree of rRNA contamination between samples could give the erroneous conclusion that m^6^A in mRNA is actually changing between experimental conditions.

We first wanted to determine if rRNA contamination is a problem when analyzing poly(A) mRNA samples. To test this, we used published RNA-Seq and m^6^A-Seq datasets in which the purified mRNA was used in mass spectrometry analysis for m^6^A and subsequent RNA-Seq and m^6^A-Seq analysis. In each of the examined datasets, the authors used either two rounds of oligo(dT) purification, or they coupled oligo(dT) purification to rRNA depletion methods^18,19^.

To determine if rRNA was present in these samples, we downloaded the raw sequencing reads from the deposited RNA-Seq and m^6^A-Seq datasets and aligned them to the human and mouse rDNA sequences to quantify rRNA-mapping reads, which are normally removed in most alignment protocols^20^. Here we found that rRNA reads accounted for a substantial fraction of total RNA-Seq and m^6^A-Seq reads, ranging from 10% to 60% in these datasets from different studies (**Fig. 1A**). The presence of rRNA in these samples highlights the difficulty in removing rRNA despite the high attention given to mRNA purification.

**Fig.1:**
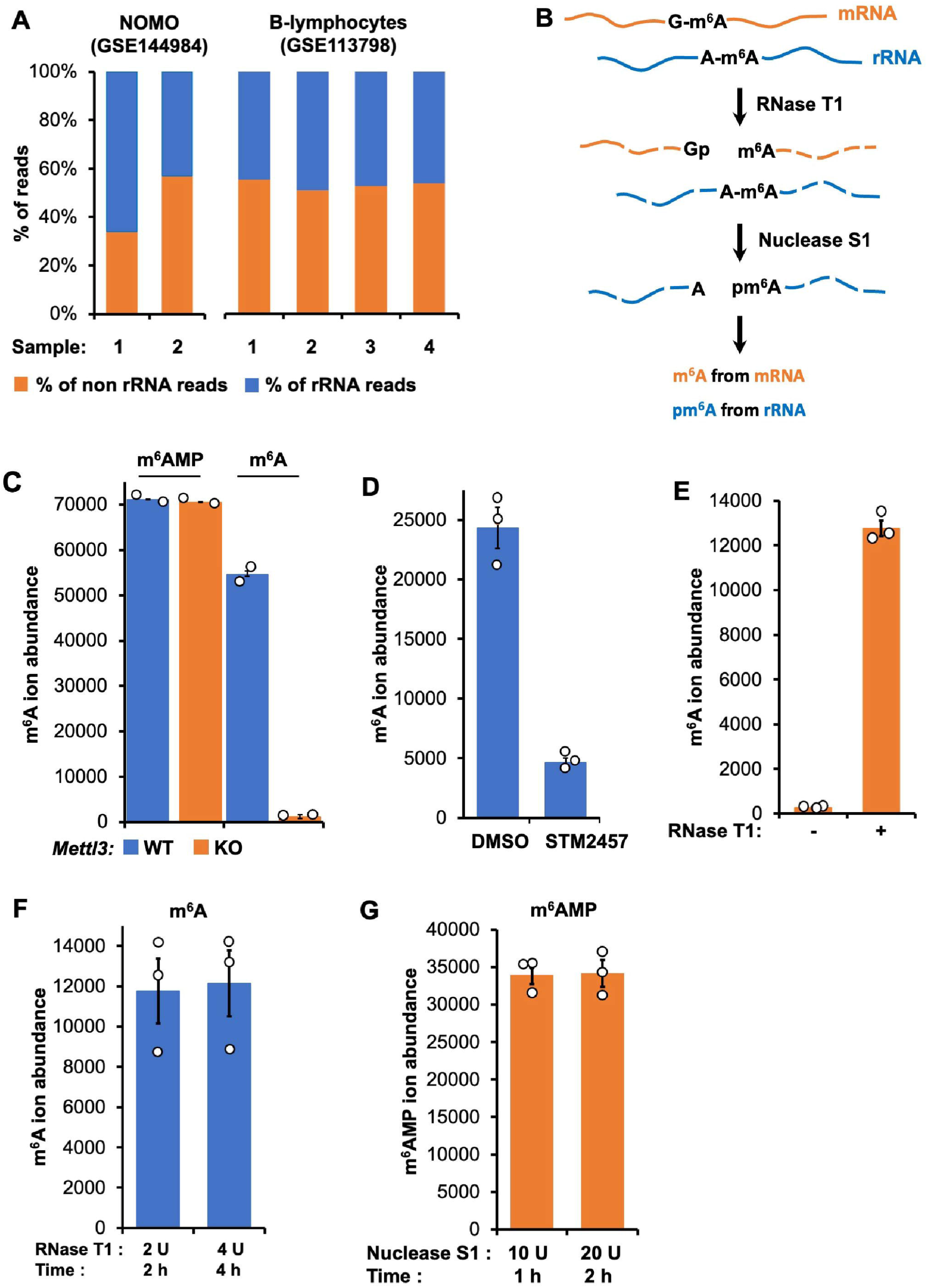
Phospho-tag m^6^A assay quantifies m^6^A in GAC sequence context in mRNA. **(A)** Contamination of rRNA in published RNA-seq and m^6^A-seq datasets in which the purified mRNA was used in both mass spectrometry analysis and subsequent RNA-seq and m^6^A-seq analysis. **(B)** Schematic showing cleavage by RNAse T1. RNase T1 specifically hydrolyzes single-stranded RNA at guanosine residues giving rise to 3’-GMP leaving a 5’OH on the adjacent nucleotide or oligoribonucleotide. Subsequently Nuclease S1 hydrolyzes RNA to nucleotide monophosphates (NMPs) with 5’-phophates and m^6^A residue is released as 5’-OH-m^6^A. **(C)** m^6^A and m^6^AMP ion abundances measured in wild-type and *Mettl3 KO* mESCs. Here m^6^A is liberated from a GAC sequence context in mRNA while m^6^AMP originates from non-GAC sequence contexts such as rRNA. **(D)** m^6^A ion abundance is markedly reduced (92%) in HEK293T cells treated with 30 μM Mettl3 inhibitor STM2457 compared to cells treated with DMSO for 6 hours. **(E)** m^6^A signal is dependent on RNase T1. When RNase T1 is omitted from the reaction the m^6^A signal is markedly diminished. **(F)** RNase T1 step is optimized. Increasing the time or the amount of RNase T1 did not further increase the yield of m^6^A. **(G)** Nuclease S1 step optimized for time of incubation and enzyme units added to the reaction. Note: The input RNA was >200nt fraction and data is based on 3 independent biological replicates and triple injections were performed for each replicate and errors bars represents standard deviation.

We next examined RNA-Seq and m^6^A-Seq datasets from experiments that reported changes in m^6^A levels in mRNA, such as a recent study which found elevated m^6^A in acute myeloid leukemia^18^ and decreased m^6^A during stress response in major depressive disorder (MDD)^19^. We again examined the RNA-Seq datasets that used poly(A) RNA prepared in the same way as the poly(A) RNA used for mass spectrometry. Here, in these datasets, which were reported to contain higher m^6^A levels^18,19^, exhibited notably lower rRNA levels in input RNA-Seq dataset (**Fig. 1A**).

Notably, we were unable to find studies where the authors addressed the possibility of rRNA contamination by measuring the rRNA levels. Thus, rRNA contamination could account for the observed “dynamic” changes in m^6^A levels in mRNA. Overall, these data highlight the difficulty in removing rRNA from mRNA samples, and the need to develop new methods that overcome this unavoidable contamination.

### Phospho-tag m^6^A assay: A method for selective detection of m^6^A mediated by METTL3/14 in total cellular RNA

Because obtaining highly pure mRNA samples is probably unrealistic for most laboratories, we wanted to develop an m^6^A quantification protocol that does not require mRNA purification. Thus this protocol must selectively measure mRNA-derived m^6^A in total RNA samples despite the vastly higher levels of rRNA and snRNA, which also contain m^6^A.

To measure m^6^A in mRNA, we took advantage of the unique sequence context of m^6^A in mRNA and other RNA polymerase II-derived transcripts. In mRNA, m^6^A is usually, but not always, preceded by a G^21,22^. This sequence preference reflects the methylation specificity of METTL3/METTL14, the heterodimeric enzyme that synthesizes m^6^A^23,24,25^. m^6^A mapping studies, along with earlier biochemical analysis, found that ∼75% of m^6^A is in a G-m^6^A-C sequence context, while 20-25% of m^6^A can be found in a A-m^6^A-C sequence context^21,22,26^. Currently, there are no known pathways that would selectively methylate m^6^A in a G-m^6^A-C context versus an A-m^6^A-C context. For this reason, m^6^A in a G-m^6^A-C context could serve as a proxy for the total level of m^6^A in the mRNA transcriptome.

Importantly, m^6^A in possible contaminating RNA is found in different sequence context. For rRNA, m^6^A is found exclusively in a A-m^6^A context^27^. In the U6 snRNA, m^6^A is found in a C-m^6^A context^28^. Therefore, any m^6^A in a G-m^6^A context reflects m^6^A from mRNA, not a contaminant.

In contrast, m^6^A in 18S and 28S rRNA is synthesized by METTL5 and ZCCHC4, respectively^29,30^, while m^6^A in U6 snRNA is catalyzed by METTL16^31^. Since only m^6^A in mRNA can be preceded by a G, the G-m^6^A sequence context can be used as a selective mark for m^6^A levels in mRNA.

To develop an assay to selectively quantify mRNA-derived m^6^A, we therefore took advantage of RNase T1, which selectively cleaves after G. Unlike other ribonucleases, RNase T1 leaves a 5’ hydroxyl after cleavage (**Fig. 1B**). Therefore, all m^6^A residues in a G-m^6^A-C sequence context are cleaved to HO-m^6^A-C-N (where N indicates one or more residues). Next, nuclease S1 is added, which cleaves all remaining phosphodiester bonds, including A-m^6^A in rRNA and C-m^6^A in snRNA. Nuclease S1 cleavage leaves a 5’ phosphate. As a result of nuclease S1, m^6^A residues in rRNA and snRNA contain a 5’ phosphate, i.e., m^6^AMP (**Fig. 1B**). However, cleavage of the RNase T1 digestion product OH-m^6^A-C-N by nuclease S1 liberates the non-phosphorylated m^6^A.

Overall, this “phospho-tag” assay is expected to generate two forms of m^6^A, i.e., m^6^A with a 5’-hydroxyl or a 5’ phosphate (referred to henceforth as m^6^A and m^6^AMP). Thus, the moiety on the 5’ of m^6^A, either hydroxyl or phosphate, provides a simple and selective way to determine if the m^6^A derives from mRNA, or contaminating rRNA or snRNA.

### The phospho-tag m^6^A assay is selective only for m^6^A generated by METTL3/14, not ZCCHC4, METTL5, or METTL16

We first asked if the phospho-tag assay only detects m^6^A generated by METTL3/METTL14. We therefore prepared total RNA from wild-type mouse embryonic stem (mES) cells as well as *Mettl3* knockout mES cells^32^. We used the *Mettl3* knockout mES cells generated by Hanna and colleagues, which exhibit a >99% reduction in m^6^A levels in highly purified mRNA^32,33^. Thus, *Mettl3* knockout mES cells are an ideal system to establish whether any of the m^6^A signal in the phospho-tag assay derives from mRNA.

Using the phospho-tag assay, m^6^A was readily detectable in total RNA prepared from wild-type mES cells (**Fig. 1C**). In contrast, the m^6^A signal was reduced by 97.1% in total RNA prepared from *Mettl3* knockout ES cells (**Fig. 1C**). Notably, m^6^AMP levels were high in both wild-type of *Mettl3* knockout samples (**Fig. 1C**), consistent with the idea that *Mettl3* knockout does not affect m^6^A in rRNA or snRNA. In addition, we treated HEK293T cells for 6 h with 30 μM STM2457, a selective METTL3 inhibitor^34^. Here we also found that m^6^A levels were markedly reduced in total RNA (**Fig. 1D**), further supporting the idea that the phospho-tag assay selectively measures m^6^A generated by METTL3/METTL14 and not other sources of m^6^A.

We next confirmed that the m^6^A signal in the phospho-tag assay is dependent on RNase T1 and does not reflect endogenous 5’-hydroxyl m^6^A RNAs. Upon removal of the RNase T1 step, which is required to generate m^6^A with a 5’ hydroxyl, an essentially complete abolition (98%) of the m^6^A signal was seen (**Fig. 1E**). Overall, these experiments demonstrate that the RNase T1 step is needed to generate m^6^A, and that the any detected m^6^A in the phospho-tag assay was present in a G-m^6^A context.

We next wanted to establish whether the enzymatic steps had gone to completion. To test this, we used 10 μg RNA from HEK293T cells and compared our RNase T1 treatment (2 h, 2 U enzyme, 37°C) to treatments associated with higher degrees of RNase T1 activity: 4 h reaction times, and adding additional RNase T1 (an additional 2 U added after 2 h, followed by an additional 2 h incubation). In both cases, we found no further increase in m^6^A levels (**Fig. 1F**).

We also asked if the nuclease S1 was also optimized. With nuclease S1, the standard reaction condition is 1 h, 10 U enzyme, at 37°C. We tested both extending the reaction time or adding more enzyme. In neither case did we see any increase in the total amount of nucleotide monophosphate levels (**Fig. 1G**). Based on these results, the RNase T1 and nuclease S1 steps were considered optimized for input RNA levels up to 10 μg.

Overall, these studies demonstrate an assay for selective detection of METTL3/METTL14-derived m^6^A in total RNA samples despite the presence of vastly larger amounts of rRNA- and snRNA-derived m^6^A.

### A phospho-tag assay for cap m^7^G to quantify mRNA transcript levels

In the traditional m^6^A quantification assay, m^6^A levels are normalized to total adenosine levels in the sample^8^. However, this approach is problematic since total adenosine levels are influenced by the levels of contaminating rRNA/snRNA, as described above, and the lengths of poly(A) tails. Poly(A) tails is particularly problematic in the m^6^A field since m^6^A recruits deadenylases to mediate mRNA degradation^35,36^. Therefore, adenosine is problematic for normalizing the m^6^A levels.

Therefore, we decided to develop a new normalization strategy. Rather than normalizing m^6^A to A, we decided to normalize m^6^A levels to transcript copy levels. A unique feature of each RNA polymerase II transcript, but not rRNA or snRNA, is the presence of an m^7^G cap^37^. This “cap m^7^G” comprises m^7^G followed by a triphosphate bridge to the first transcribed nucleotide^37^. In addition to cap m^7^G, “internal m^7^G” is also found in other classes of RNAs, including mRNA, rRNA and tRNA^38,39,40^ Therefore, the phospho-tag assay needs to distinguish cap m^7^G from internal m^7^G.

To distinguish cap m^7^G from internal m^7^G, we developed a second approach that again uses phosphate to mark the origin of m7G (**Fig. 2A**). These steps are designed to occur in the same tube used for m^6^A quantification above. In this way, parallel sample handling is avoided, thus reducing variability. As a result, m^6^A and cap m^7^G can be quantified from a one-pot reaction.

**Fig. 2:**
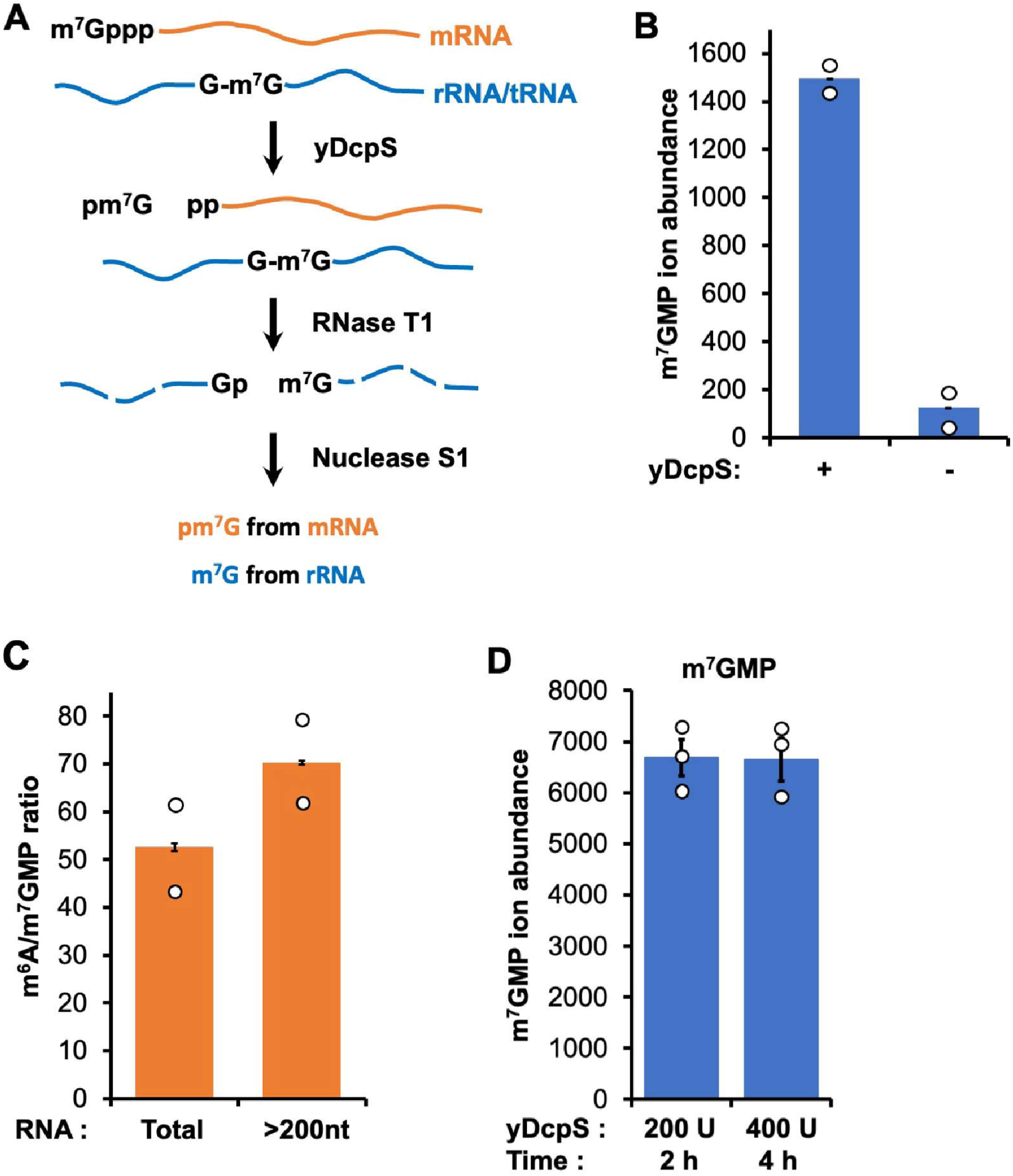
m^7^GMP signal is derived from the cap m^7^G. **(A)** yDcpS catalyzed decapping of mRNA liberates cap m^7^G as m^7^GMP and internal m^7^G in mRNA which predominantly exists in a GGN sequence context is liberated as 5’-OH-m^7^GN after cleavage by RNase T1 and subsequently released as 5’-OH-m^7^G after Nuclease S1 digestion. **(B)** m^7^GMP is primarily derived from the cap m^7^G since omission (-) of yDcpS resulted in a marked reduction (91.1%) of the m^7^GMP signal. The residual non-cap m^7^GMP likely derives from the internal m^7^G nucleotides in cellular RNAs that are not preceded by G, snice these will be liberated as m^7^GMP after the nuclease S1 step. Average m^7^GMP ion abundances in total RNA from wild-type mES cells. **(C)** The m^6^A:m^7^GMP ratio is increased in > 200nt RNA fraction compared to the total RNA. **(D)** yDcpS decapping step is optimized. Increasing the time or the amount of yDcpS did not further increase the yield of m^7^GMP. Note: The data is based on 3 independent biological replicates and triple injections were performed for each replicate and errors bars represents standard deviation.

In order to mark cap m^7^G from internal m^7^G, we developed a protocol that takes advantage of both RNase T1 and the decapping enzyme yDcpS^41^. Internal m^7^G is usually preceded by a nonmethylated G: G-m^7^G^42,43^. As a result, after RNase T1 digestion, internal m^7^G will be released with a 5’-hydroxyl. Therefore, we wanted to mark cap m^7^G with a 5’-phosphate. We therefore used yDcpS, which releases the cap m^7^G as a m^7^GMP^41^. yDcpS is different from other decapping enzymes such as Dcp2, Nudt12, Nudt15 and RppH which release cap m^7^G as m^7^GDP^44^ (**Fig. 2A**). We could not use these other decapping enzyme since m^7^GDP contains a diphosphate, which is not readily detected on mass spectrometry due to ion suppression^45^. Therefore, yDcpS provides a unique opportunity to release m^7^G with a phosphate that can reveal that the m^7^G is derived from the cap.

We first asked if the m^7^GMP signal derives from the cap. The phospho-tag assay generated readily detectable m^7^GMP when using total RNA from wild-type mES cells (**Fig. 2B**). This was primarily derived from cap m^7^G since omitting yDcpS (2 h, 200 U, 37°C), resulted in a marked reduction (91.1%) of the m^7^GMP signal (**Fig. 2B**). The residual non-cap derived m^7^GMP likely derives from internal m^7^G nucleotides in cellular RNAs that are not preceded by G^38^, since these will generate m^7^GMP after the nuclease S1 step.

Since some m^7^GMP derives from internal m^7^G, we included a “no yDcpS” control for each sample in the following experiments. By subtracting this background signal, cap m^7^G can be more accurately determined.

One potential confounding factor when measuring cap m^7^G is that some cap m^7^G derives from short transcripts, such as enhancer RNAs^46,47^. These short transcripts may have a very low m^6^A/cap m^7^G ratio since the transcripts are too short to contain m^6^A. Since our goal is to measure m^6^A in mRNA, we used a size fractionation step. In this step, the extracted RNA is subjected to a silica-based purification column that contains wash steps that cause shorter (<200 nt) RNA to be removed.

We asked if the m^6^A/cap m^7^G ratio is different in these two samples. In the fraction containing >200 nt RNA, the m^6^A/cap m^7^G ratio was ∼25% increased (**Fig. 2C**), consistent with the idea that the inclusion of small RNAs artificially lowers the m^6^A/cap m^7^G ratio. We therefore used the >200 nt fraction for all subsequent experiments.

As before, we determined if the yDcpS treatment was optimized. For these experiments, we again used 10 μg of input RNA after >200 nt size selection from HEK293T cells. Increasing the time or the amount of enzyme did not further increase the yield of m^7^GMP (**Fig. 2D**). Thus, the overall phospho-tag assay was considered optimized for both m^6^A and m^7^GMP detection for this amount of input RNA.

### LC-MS/MS method validation

Compared to other m^6^A quantification methods^48^, our phospho-tag m^6^A assay not only selectively measured mRNA-derived m^6^A and m^7^GMP in one sample injection, but also resolved m^6^A isomers 1-methyladensosine (m^1^A) and 2’-*O*-methyladenosine (Am) detection by either LC chromatographic retention time or MS/MS fragment ions (**Supplementary Fig. S1**). m^1^A and m^6^A share the same major fragment ion (quantifier) of methyladenine (M+H, 150.07) and same MRM transition of 282.1→ 150.1, but they can be readily separated by retention time (1.8 vs 2.1 min). While Am is not abundant in our RNA digests, its detection can be distinguished from m^6^A and m1A by fragment ion of adenine (M+H, 136.06) and transition of 282.1→136.1.

We next validated the phospho-tag m^6^A assay for method sensitivity, linear range, matrix effects, precision and accuracy. The linearity for the detection of m^6^A and m^7^GMP was determined by constructing external calibration curves for unlabeled synthetic standard solution of m^6^A and m^7^GMP. To assess the extent of matrix effect on LC-MS/MS measurements, we compared calibration curves prepared from solvent blank (80% methanol, 20% H2O) and matrix solution containing RNA digestion assay buffers (**Fig. 3A**). In the absence of spiked-in internal standards, both m^6^A and m^7^GMP measurements showed fair linearities (R2>0.969) within a dynamic linear range of up to 1μm analytes. The presence of matrix (i.e., RNA digestion buffer) resulted 17% and 26 % decrease in m^6^A and m^7^GMP detection sensitivity, respectively.

**Fig. 3:**
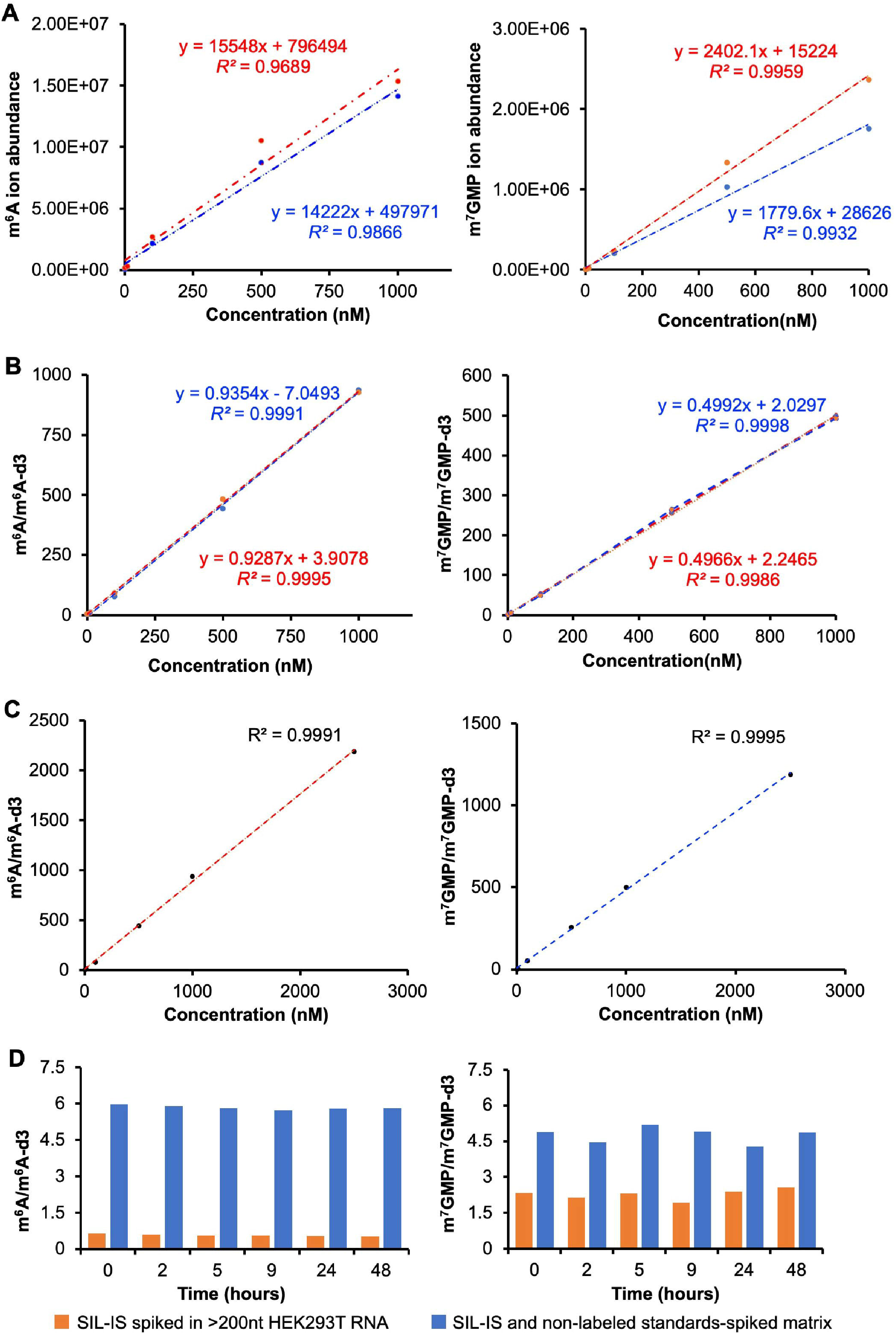
Phospho-tag m^6^A assay linearity is improved by use of SIL-IS. **(A)** Comparison of calibration curves prepared either in solvent blank (80% methanol, 20% H2O) and matrix solution (Phospho-tag assay digestion buffers). In the absence of spiked-in SIL-IS, matrix effect resulted in 17% and 26 % decrease in m^6^A and m^7^GMP detection sensitivity, respectively. **(B-C)** SIL-IS eliminated matrix effect and improved the linearities of both m^6^A and m^7^GMP. **(D)** Stability of m^6^A and m^7^GMP. Unlabeled m^6^A and m^7^GMP standards and SIL-IS spiked into matrix and SIL-IS spiked into RNA digestion mixture and analyzed over a time span of 48 hours. Ion abundance ratio relative to SIL-IS shows overall constant readout for both analytes over 48h at 4°C (**Fig. 3D**). Note: All the data is based on triple injections of each sample.

To eliminate the observed matrix effect and to correct day-to-day, batch-to-batch variability in LC-MS/MS measurement, we spiked in stable isotope-labeled internal standard (SIL-IS), m^6^A-d3 and m^7^GMP-d3, into sample matrix at a constant concentration of 1 nM and 5 nM, respectively. As expected, SIL-IS not only eliminated matrix effect, but also improved the linearities of both m^6^A and m^7^GMP calibration curves to R^2^ greater than 0.999 (**Fig. 3B)**. The linear range for both analyte detections were also extended to 2 μm (**Fig. 3C)**. The limit of detection (LOD) and the limit of quantification (LOQ) for m^6^A and m^7^GMP calculated as 3 and 10 times blank standard deviation to slope ratio (3*STDEV/slope, 10*STDEV/slope)) were as follows: LOD 0.03 nM and LOQ 0.1 nm for m^6^A, and LOD of 0.3 nM and LOQ of 1.1 nm for m^7^GMP.

Several factors can affect the accuracy of measurement including extraction efficiency, stability of the analyte, adequacy of the chromatographic separation and the purity of reference standards^49^. To test the robustness of phospho-tag m^6^A assay, we measured the coefficient of variation (CV) in repeated injection of 12 RNA samples and recovery rate of spiked standards to further validate the method precision and accuracy. The recovery rates for 5 nM and 10 Nm m^6^A and m^7^GMP standard spiked into RNA digests were in the range of 88-116% which is well within the accepted range of 80-120%. We then assessed the stability of m^6^A and m^7^GMP chemical standards spiked into RNA-free matrix and m^6^A and m^7^GMP derived from RNA digestion mixture over a time span of 48h. We observed an overall constant readout of ion abundance ratio relative to deuterated internal standard for both analytes over 48h at 4°C, with the coefficient of variations for RNA sample and chemical standard well within 9.7% (**Fig. 3D**). Note that the total LC-MS/MS sample run is 8 min, phospho-tag can easily achieve >300 sample run with accuracy and robustness.

### Input requirements for the phospho-tag m^6^A assay

We next determined the minimal RNA input requirements needed to quantify m^7^GMP and m^6^A. Our optimizations for RNase T1 and yDcpS used 10 μg, but it would be desirable to know the minimum amount of input RNA that would provide accurate m^6^A and cap m^7^G measurements. We tested 1, 2.5, 5, 7.5 and 10 μg amounts of input RNA using the phospho-tag assay in two biological replicates. At all these RNA sample inputs, we were readily able to measure m^6^A and m^7^GMP (**Fig. 4A**). Over this range of RNA input, the m^6^A and m^7^G levels showed linearity (correlation coefficients r^2^□> □ 0.995). These data suggest that m^6^A and m^7^GMP measured using RNA input levels as low as 1 μg can be reliably measured in this assay.

**Fig. 4:**
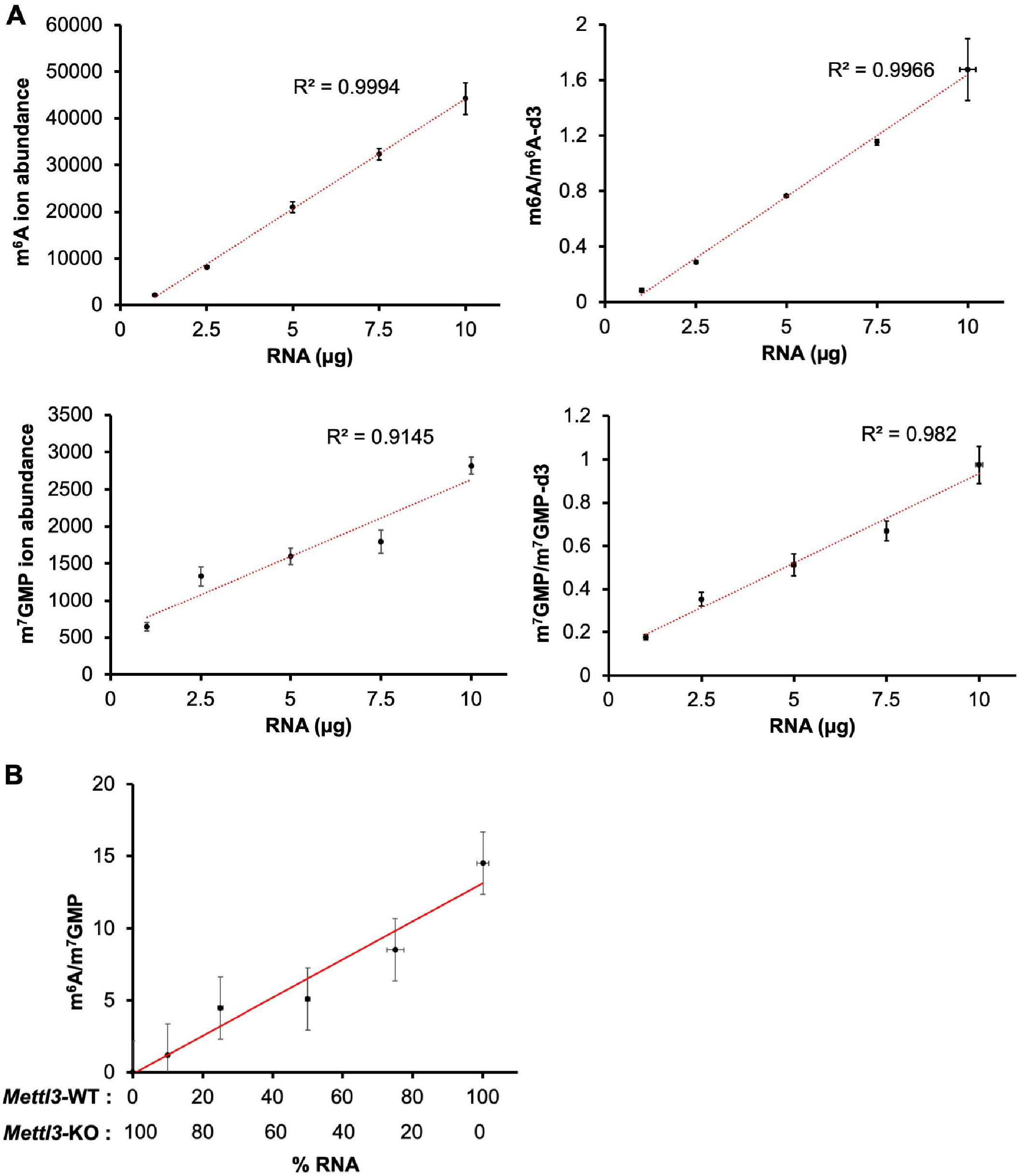
Quantitative accuracy of phospho-tag m^6^A assay for assessment of m^6^A and m^7^GMP. **(A)** Linearity was observed in m^6^A levels and m^6^A/m^6^A-d3 levels (R^2^>0.995) within range 0-10 μg of >200nt RNA input. m^7^GMP showed poor linearity (R^2^=0.914) with increasing RNA input amounts, however this was markedly improved (R^2^=0.982) with m^7^GMP/m^7^GMP-d3. **(B)** Phospho-tag assay shows quantitative accuracy. >200nt RNA from wild-type and *Mettl3* knockout mES cells mixed in specific ratios and estimated the quantified the m^6^A:m^7^GMP ratio. The m^6^A:m^7^GMP ratio directly correlated with the fraction of wildtype RNA mixed with the *Mettl3* knockout RNA.

Notably, the limiting factor for accurate quantification of the m^6^A:m^7^GMP ratio in the phospho-tag m^6^A assay was the detection of m^7^GMP. The m^6^A measurements were linearly increased with increase in the RNA input from 1 to10 μg (r^2^□> □ 0.995) (**Fig. 4A**).

However, the m^7^GMP measurements didn’t display the degree of linearity with increase in RNA input as compared to m^6^A (r^2^□ = □0.91). The linearity of m^7^GMP was markedly improved (r^2^□ = 0.98) by use of m^7^GMP-d3 spike in at constant concentration of 5 nM in each sample post methanol precipitation step. The poor detection of m^7^GMP with low input RNA likely reflects the reduced ionization efficiency of m^7^GMP, a phenomenon seen with other phosphorylated molecules^45^.

We next assessed the quantitative accuracy of the phospho-tag m^6^A assay. To do this, we prepared total RNA from wild-type and *Mettl3* knockout mES cells. We then mixed these RNAs in specific ratios and estimated the quantified the m^6^A:m^7^GMP ratio. For all experiments, the sum of wild-type and *Mettl3* knockout RNA was 10 μg. Here we found that the m^6^A:m^7^GMP using the phospho-tag m^6^A assay directly correlated with the fraction of wildtype RNA mixed with the *Mettl3* knockout RNA (**Fig. 4B**). Overall, these studies suggest that the phospho-tag m^6^A assay shows high quantitative accuracy.

### Quantitative analysis of m^6^A/m^7^GMP ratios using stable isotopically labeled-internal standards (SIL-IS)

To increase the quantitative accuracy of m^6^A and m^7^GMP detection, we analyzed samples with stable isotopically labeled-internal standard (SIL-IS) of m^6^A and m^7^GMP. Use of SIL-IS improves reproducibility between injections, compensates for the loss of sensitivity during a batch run of samples, and accounts for matrix effects that can happen during the ionization process^50,51^.

To test this, we prepared input RNA from HEK293T cells and measured the m^6^A:m^7^GMP ratio in three biological replicates in which we either did not add standards or added standards. For the samples with standards, we mixed the m^7^GMP-d3 and m^6^A-d3 standards after the methanol precipitation step at a concentration of 5 nM and 1 nM, respectively. We then calculated the m^6^A:m^7^GMP ratio for both sets of samples. In the case of the samples without standards the average m^6^A:m^7^GMP ratio was 20.3 +/- 10.3 % RSD, while the average m^6^A:m^7^GMP ratio was 1.7 +/- 3.9% when standards were used (**Fig. 4C**). These results show that the variability in measurement is markedly reduced when using the SIL-IS standards.

An additional benefit of using SIL-IS is that the m^6^A:m^7^GMP ratio can be determined using exact amounts of m^6^A and m^7^GMP. Thus, results with SIL-IS also reveal the exact ratio of m^6^A per transcript. For all subsequent experiments SIL-IS was used to quantify the m^6^A:m^7^GMP ratio.

## DISCUSSION

A major problem with m^6^A assays is that they can be highly affected by the way in which the sample is prepared. Most assays rely on poly(A) purification, but this method typically is associated with residual contaminating rRNA. Additionally, poly(A) purification is associated with preferential accumulation of mRNA from 3’ ends, a region which is known to be enriched in m^6^A. Since these problems are hard to control between samples, sample-to-sample variation in m^6^A levels, as measured by mass spectrometry or other methods, may simply reflect variation in these sources of error. The phospho-tag m^6^A assay overcomes this problem by selectively measuring only m^6^A from mRNA. Thus any rRNA contamination does not contribute to the overall m^6^A measurement. Additionally, since total mRNA is used, there is no bias for 3’ ends of mRNA. As a result, all parts of the mRNA are used in this assay. Notably, the phospho-tag m^6^A assay also includes normalization to m^7^G caps, thus providing insights into the overall level of m^6^A per transcript.

The core concept of the phospho-tag m^6^A assay is that phosphates are used as an endogenously derived mass tag to indicate the origin of m^6^A. m^6^A from rRNA or snRNA will produce m^6^A with a 5’ phosphate. In contrast, m^6^A derived from mRNA will produce m^6^A as a nucleoside, i.e., with no phosphate, as long as the m^6^A was preceded by G. Importantly, ∼75% of all m^6^A in mRNA is preceded by a G^21,22,26,52,53^. Thus, the amount of m^6^A calculated per transcript can be adjusted to take into account that the value calculated in the phospho-tag assay is ∼75% of the total level of m^6^A in mRNAs. The presence or absence of phosphate changes the mobility of m^6^A in the LC step of the MS analysis and provides a unique mass. In this way, m^6^AMP cannot be accidentally measured and used in the calculation of the overall m^6^A level in mRNA.

The phosphate tag on m^6^A derived from rRNA and snRNA, but not mRNA, is a result of the selectivity of RNase T1. RNase T1 cleaves RNA after G residues, and importantly, leaves the phosphate on the 3’ end of the G. As a result, the subsequent nucleotide has a 5’ hydroxyl. Since m^6^A in mRNA typically is preceded by a G, it will contain a 5’ hydroxyl after RNase T1 treatment. In contrast, the A-m^6^A bond in rRNA and the C-m^6^A bond in snRNA will not be cleaved by RNase T1. Instead, these bonds are cleaved by nuclease S1 in the subsequent step. Nuclease S1 cleaves the RNA, leaving the phosphate on the m^6^A. In this way, the phosphate acts as a tag that reveals the RNA origin of the m^6^A.

The second core idea of the phospho-tag assay is that m^6^A levels are normalized to mRNA abundance by using the m^7^G cap as a surrogate for the overall mRNA transcript levels. In previous studies, m^6^A was normalized to total A levels in a sample. This is problematic because rRNA is typically present in mRNA samples, and thus, this normalization is highly affected by the level of rRNA contamination. In contrast, since rRNA and snRNA lack m^7^G caps, their presence does not affect the overall quantification of m^6^A/m^7^GMP.

Since m^6^A is normalized to transcript levels, it is important to keep in mind that changes in m^6^A/m^7^GMP levels might reflect changes in mRNA 3’UTR lengths. In some conditions, 3’UTR lengths change as a result of regulated polyadenylation site selection. Thus, if an mRNA becomes longer due to a longer 3’UTR, it may have more m^6^A per transcript. However, this does not mean that the stoichiometry of m^6^A sites has increased. Thus, any observed changes in m^6^A levels between two samples should be orthogonally validated using site-specific measurements of m^6^A stoichiometry using assays such as SCARLET^54^.

The cap-derived m^7^G is also selectively marked by a phosphate. This is achieved using yDcpS, an enzyme the selectively hydrolyzes m^7^G that is part of the mRNA cap. yDcpS releases m^7^G with a 5’ monophosphate. Importantly, this reaction is performed along with the RNase T1 reaction in a single pot. As a result, internal m^7^G nucleotides, which are typically found in rRNA and tRNA preceded by a G, are released from RNA as a m^7^G with a 5’ hydroxyl. In this way, the cap-derived m^7^G can be selectively measured based on its unique mass and retention time conferred by its 5’ phosphate.

Notably, there is a small amount of m^7^G in cellular RNA that is not preceded by a G. This is seen in samples that are not treated with yDcpS. Therefore, a “no yDcpS” control should be used to establish this background level of m^7^GMP. In contrast, very little m6A containing a 5’ hydroxyl is generated in samples where RNase T1 was omitted. Nevertheless, a “no RNase T1” control is important to ensure that the m6A signal indeed derives from cellular mRNA.

The phospho-tag m^6^A assay was validated using a *Mettl3* knockout embryonic stem cell line. Importantly, total RNA was used in these experiments. Thus, sample preparation was highly simplified. Despite the large amount of m^6^A from rRNA in these samples, no m^6^A was detected since this assay is highly selective for m^6^A derived from mRNA.

Overall, we expect that the phospho-tag m^6^A assay will allow rapid and simple measurements of m^6^A from essentially and RNA sample without the need for tedious poly(A) purification. Also, since this reaction is a single-pot reaction, it can be used in medium- and high-throughput assays for m^6^A measurements. We expect that the phospho-tag m^6^A assay will reveal whether m^6^A is dynamic and will help to identify the specific signaling or disease contexts in which these dynamics occur.

## MATERIALS AND METHODS

### rRNA mapping

Publicly available raw RNA-Seq and m^6^A-Seq reads from Gene Expression Omnibus database (GEO) were downloaded from two studies (GEO accession: GSE144984 and GSE113798). The fastq files quality was checked with MultiQC tool^55^. Adapters were removed using Trimmomatic^56^. The adapter-trimmed reads were mapped to the human rDNA complete unit (KY962518.1) using Bowtie2^57^ (version 2.4.2) with --local option. For each sample number of mapped and unmapped reads were used to calculate the percentage of rRNA reads. The rRNA reads from biological or technical replicates were averaged.

### Cell line culture

HEK293T/17 and mouse embryonic fibroblast cells (mEFCs) were cultured in 1× DMEM (Life Technologies #11995-065) with 10% FBS, 100□U□ml^−1^ penicillin and 100□μg□ml^−1^ of streptomycin under standard tissue culture conditions. Cells were split using TrypLE Express (Life Technologies) according to manufacturer’s instructions. Cells were harvested after reaching 80% confluency. Mouse embryonic stem (mESCs) were previously described by Geula et al (^32^),and were a kind gift from S. Geula and J.H. Hanna (Weizmann Institute of Science). All mESCs were grown in tissue culture plates precoated with 0.1% gelatin (EmbryoMax ES-006-B) in mESC media (KnockOut DMEM (Gibco #10829018), 15% heat-inactivated fetal bovine serum (FBS) (Gibco #26140079), 100 μg/ml streptomycin (Gibco #15140122), 100 U/ml penicillin, 1x GlutaMax (Gibco #35050061), 55 μM β-mercaptoethanol (Gibco #21985023), 1x MEM non-essential amino acids (Gibco #11140076), 3 μM CHIR99201 (Sigma Aldrich SML1046), 1 μM PD0325901 (APExBIO # A3013), 1000 U/ml LIF (Millipore # ESG1107).

### RNA isolation

RNA was isolated using TRIzol™ LS Reagent (ThermoFisher #10296010) following manufacturer’s instructions except after adding chloroform the sample was transferred to pre-spined MaXtract High Density (Qiagen #129065) and centrifuged at 12,000*g* for 15 mins at 4°C to phase sperate the aqueous phase from the organic phase. Next the aqueous phase containing RNA was transferred into a fresh tube and standard protocol was followed for total RNA isolation. Next, the isolated RNA was treated with DNase I (Invitrogen™ #18047019) at 37°C for 1 hour to remove traces of contaminating DNA. The DNase I treated RNA was further purified into small (<200nt) and large (>200nt) RNA fractions using RNA Clean & Concentrator™-25 (Zymo Research # R1017) following the manufacturer’s instructions. The large (>200nt) RNA fraction was used as the input for Phospho-tag assay.

### Enzymatic digestion of RNA

10 μg of >200nt RNA (in nuclease-free water) was decapped using 200 U yDcpS and yDcpS buffer (NEB, #M0463S) at 37°C in a thermomixer at 800rpm pulse shaking. Next 2 U of RNase T1 (INVITROGEN #AM2283) was added to the tube and incubated at 37°C in a thermomixer at 800rpm pulse shaking. To the above tube,10 U of S1 Nuclease (INVITROGEN #18001-016) was added, and sample was incubated for 1 hour at 37°C in a thermomixer at 800 rpm pulse shaking to digest RNA to mononucleotides. Next 4 volumes of 100% methanol was added to facilitate precipitation of all enzymes present in the hydrolysate before the sample injection. The samples were centrifuged at 16,000*g* for 30 min at 4°C and supernatant was carefully transferred into a fresh 1.5 mL tube without disturbing the pellet. Additionally, for each phospho-tag m^6^A assay two control reactions were step up: (i) yDcpS is omitted (ii) RNase T1 is omitted. The signals from these two controls are subtracted from the sample in which all enzymes were added.

### Preparation of calibration with and without SIL-IS

For preparation of calibration solutions synthetic standard of N6-methyladenosine (m^6^A) was purchased from Selleck Chemicals, Houston, Texas, USA (Catalog No. S3190) and 7-Methyl-guanosine-5’-monophosphate (m^7^GMP) was purchased from Jena Bioscience, Jena, Germany (Catalog No. NU-1135S). N1-methyadenosine (m1A) was purchased from (Carbosynth #NM03697). The deuterated standards m^6^A-d3 and m^7^GMP-d3 were custom synthesized from Toronto Research Chemicals, Ontario, Canada).

Synthetic standards of m^6^A and m^7^GMP were dissolved in pure water at a final concentration of 1 mM each. From these stocks two set of calibration solutions were prepared in the range of 0 nM to 1 μM in sample matrix and 80% methanol. The sample matrix consists of all the reaction and enzyme storage buffers in 80% methanol. Additionally, calibration sets were also spiked with SIL-IS, m^6^A-d3 and m^7^GMP-d3 at constant a final concentration of 1 nM and 5 nM throughout all calibration samples, respectively to correct for sample matrix effect. For each Phospho-tag assay, 50 μl of methanol precipitated hydrolysate was aliquoted into HPLC vials and m^7^GMP-d3 and m^6^A-d3 was spiked in at a final concentration of 5 nM and 1 nM, respectively.

### Stability test for m^6^A and m^7^GMP

To assess the stability of m^6^A and m^7^GMP we performed repeated injections of two set of samples spiked with synthetic m^6^A, m^7^GMP and SIL-IS m^6^A-d3 and m^7^GMP-d3 at 10 nM over a period of 48 hours in sample matrix and 80% methanol. We performed another set of repeated injections of digested RNA(>200nt) spiked with SIL-IS at 10 nM over 48 hour period. We then calculated % RSD (relative standard deviation) for all three sample sets. The samples were maintained at 4°C.

### Procedure for LC-MS/MS measurements

RNA digests were analyzed by LC-MS/MS using a platform comprised of an Agilent Model 1290 Infinity II liquid chromatography system coupled to an Agilent 6460 Triple Quadrupole mass spectrometer equipped with Agilent Jet Stream Technology. Chromatography of metabolites utilized reversed-phase chromatography on a Infinity Lab Poroshell 120 EC-C18 column (Agilent Part Number:695975-902). Mobile phases consisted of: (A) 99% water, 1% acetonitrile containing 1 mM ammonium formate and 0.1% formic acid, and (B) 99% acetonitrile, 1% water containing 1 mM ammonium formate and 0.1% formic acid. The following gradient was applied at the flow rate of 0.9 ml/min: 0-0.5 min, 90% A,10% B; 0.5-1.0 min, 70% A, 30% B; 1-4 min, 30%, 70% B; 4-4.1 min, 1% A, 99% B, to 6 min, followed by a re-equilibration at 90% A for 2min. The column compartment temperature was at 25°C. The injection volume is 2 μl. MRM data were acquired in positive ion mode. Fragmentor, collision energy, and other source parameters were optimized for both quantifier and qualifier ions using Agilent MassHunter Optimizer software (version 6.0). Source parameters for m^6^A measurement were as follows: gas temperature, 230°C; gas flow, 8 l/min; nebulizer, 35 psi; sheath gas temperature, 400°C; sheath gas flow, 12 l/min; capillary voltage, 2500v; delta EMV, 400 v. The source parameters for m^7^GMP are the same as m^6^A. The quantification of m^6^A and m^7^GMP were achieved using MRM transitions for both quantifier and qualifier ions shown in Table 1.

**Table 1:** MRM transitions and source parameters selected for the analysis of m^6^A, m^6^A-d3

| Compound | RT (min) | Fragmentor (V) | Collision energy (eV) | MRM transitions (m/z) |
| --- | --- | --- | --- | --- |
| m<sup>6</sup>A | 2.05 | 109 | **18** | **282.1 > 150.1** |
|  |  |  | 54 | 282.1 > 123.0 |
|  |  |  | 50 | 282.1 > 94.0 |
| m<sup>7</sup>GMP | 1.8 | 81 | **18** | **378.08 > 166.0** |
|  |  |  | 60 | 378.08 > 148.9 |
|  |  |  | 60 | 378.08 > 123.9 |
| m<sup>6</sup>A-d3 | 2.05 | 109 | **18** | **285.1 > 153.1** |
|  |  |  | 54 | 285.1 > 126.0 |
|  |  |  | 50 | 285.1 > 96.0 |
| m<sup>7</sup>GMP-d3 | 1.8 | 81 | **18** | **381.09 > 169.09** |
|  |  |  | 60 | 381.09 > 152.06 |
|  |  |  | 60 | 381.09 > 148.9 |

The LC-MS/MS data was analyzed using Agilent MassHunter Quantitative Analysis (for QQQ).

## AUTHOR CONTRIBUTIONS

S.R.J. and A.H.M. conceived and designed the experiments. A.H.M. carried out experiments, analyzed data, and prepared figures. Q.C. developed the LC-MS/MS part of the assay and N.A. performed LC-MS/MS measurements. S.R.J. and A.H.M. wrote the manuscript with help from all the authors.

## ACKNOWLEDGEMENTS

We thank members of the Jaffrey Lab for helpful comments and suggestions. This work was supported by NIH grant R35NS111631 to S.R.J, and T32CA062948 to A.H.M.

---

**Fig-S1:**
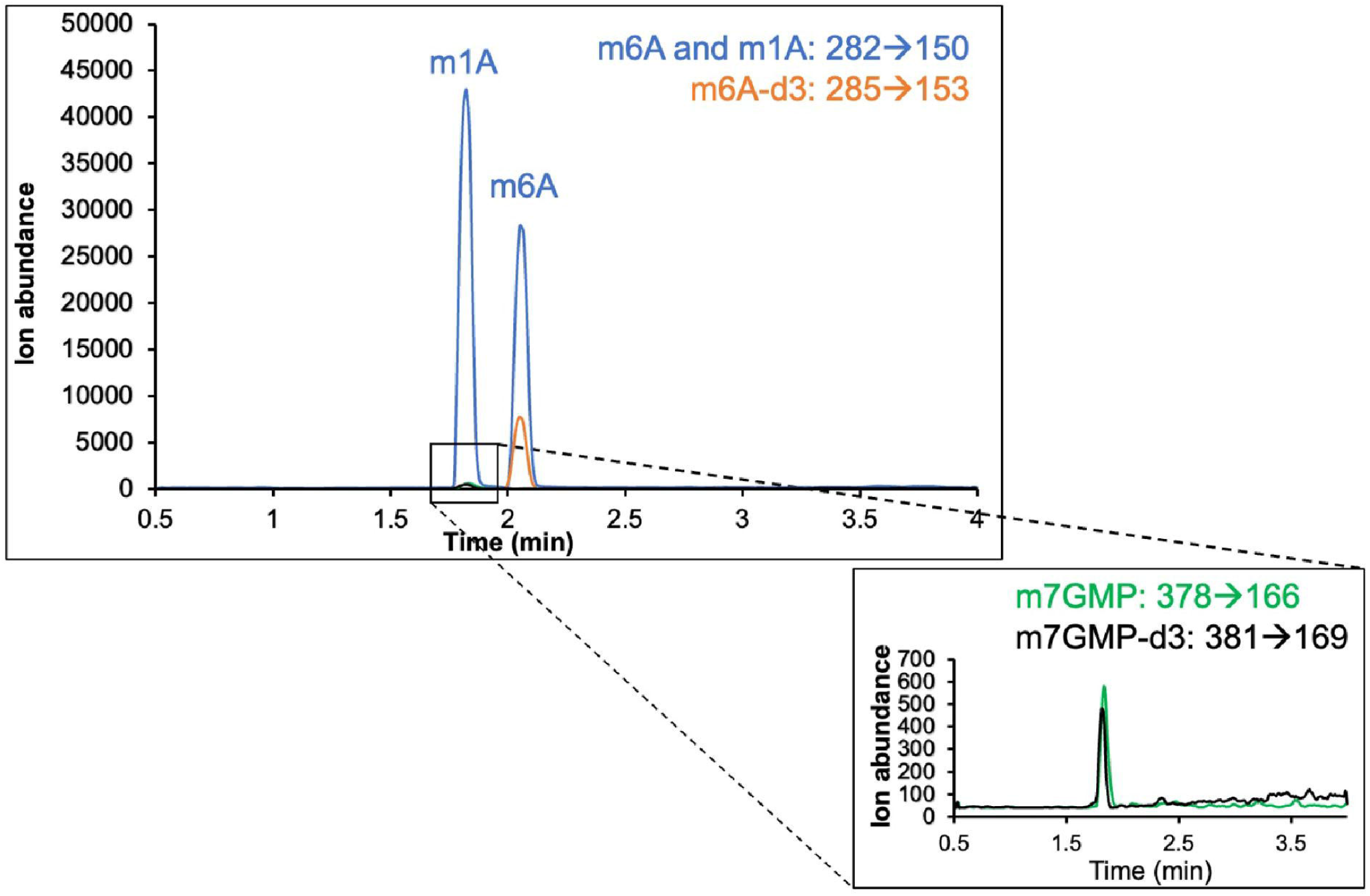
Chromatogram of m^6^A, m^6^A-d3, m^1^A, m^7^GMP and m^7^GMP-d3.

